# DNA Thermo-Protection Facilitates Whole Genome Sequencing of Mycobacteria Direct from Clinical Samples by the Nanopore Platform

**DOI:** 10.1101/2020.04.05.026864

**Authors:** Sophie George, Yifei Xu, Gillian Rodger, Marcus Morgan, Nicholas D. Sanderson, Sarah J. Hoosdally, Samantha Thulborn, Esther Robinson, Priti Rathod, A. Sarah Walker, Timothy E. A. Peto, Derrick W. Crook, Kate E. Dingle

## Abstract

*Mycobacterium tuberculosis* (MTB) is the leading cause of death from bacterial infection. Improved rapid diagnosis and antimicrobial resistance determination, such as by whole genome sequencing, are required. Our aim was to develop a simple, low-cost method of preparing DNA for Oxford Nanopore Technologies (ONT) sequencing direct from MTB positive clinical samples (without culture). Simultaneous sputum liquefaction, bacteria heat-inactivation (99°C/30min) and enrichment for Mycobacteria DNA was achieved using an equal volume of thermo-protection buffer (4M KCl, 0.05M HEPES buffer pH7.5, 0.1% DTT). The buffer emulated intracellular conditions found in hyperthermophiles, thus protecting DNA from rapid thermo-degradation, which renders it a poor template for sequencing. Initial validation employed Mycobacteria DNA (extracted or intracellular). Next, mock clinical samples (infection-negative human sputum spiked 0-10^5^ BCG cells/ml) underwent liquefaction in thermo-protection buffer and heat-inactivation. DNA was extracted and sequenced. Human DNA degraded faster than Mycobacteria DNA, resulting in target enrichment. Four replicate experiments each demonstrated detection at 10^1^ BCG cells/ml, with 31-59 MTB complex reads. Maximal genome coverage (>97% at 5x-depth) was achieved at 10^4^ BCG cells/ml; >91% coverage (1x depth) at 10^3^ BCG cells/ml. Final validation employed MTB positive clinical samples (n=20), revealed initial sample volumes ≥1ml typically yielded higher mean depth of MTB genome coverage, the overall range 0.55-81.02. A mean depth of 3 gave >96% one-fold TB genome coverage (in 15/20 clinical samples). A mean depth of 15 achieved >99% five-fold genome coverage (in 9/20 clinical samples). In summary, direct-from-sample sequencing of MTB genomes was facilitated by a low cost thermo-protection buffer.

## INTRODUCTION

*Mycobacterim tuberculosis* is the leading bacterial cause of death from infection, the World Health Organization (WHO) estimating that 10 million new tuberculosis (TB) cases and 1.2 million deaths occurred worldwide in 2018 (World Health Organization Global Tuberculosis Report 2019; https://www.who.int/tb/publications/global_report/en/). In addition, 5-10% of an estimated 1.7 billion people with latent TB infections are at risk of progressing to active disease. The greatest burden occurs in under-resourced regions of South-East Asia, Africa and the Western Pacific. There are large discrepancies between the estimated annual number of new cases (10 million) and the number reported (7 million) (WHO report https://www.who.int/tb/publications/global_report/en/). Consequently, diagnostic methods for use at the point of care, to identify ‘missing’ cases are a global priority (1, 2). Rapid diagnosis and antimicrobial resistance determination are essential to ensure appropriate TB treatment and control, particularly in light of increasing drug resistance (3, 4). In 2018, approximately 500,000 cases of rifampicin-resistant TB were identified, 78% of which were also isoniazid resistant (multi-drug resistant) (https://www.who.int/tb/publications/global_report/en/).

The application of DNA sequencing to TB molecular diagnostics yields clinically valuable information. Its utility increases with the proportion of genome obtained; from detection, to speciation and antimicrobial resistance prediction, to phylogenetic and evolutionary insights. This allows whole genome sequencing (WGS) to out-perform other rapid molecular methods (such as GeneXpert MTB/RIF, Cepheid, Solna, Sweden) due to susceptibility predictions to multiple drugs, and its combination with classical epidemiological methods which informs transmission (5–8).

WGS of MTB from early positive cultures offers markedly faster results than traditional culture-based methods which take ≤80 days. The national implementation of Illumina sequencing of Mycobacteria from early positive cultures in England provides WGS in three to four weeks, together with antimicrobial resistance predictions (5, 9). Time to WGS could be reduced further if routine sequencing could be performed direct-from-sample. Furthermore, most DNA sequencing platforms (eg Illumina) have high capital costs, require reliable power supplies, cold chain reagent shipping and highly trained staff. In contrast, the Oxford Nanopore Technologies platform, (Oxford, UK), determines nucleotide sequences via a compact, portable device powered using a laptop USB port, which can be operated in varied and challenging locations (10–13). This offers a potential direct-from-sample sequencing platform for settings with the highest burden of TB – following the third pillar of the WHO End TB Strategy, ‘intensified research and innovation’ (1). However, multiple sample-preparation issues remain to be solved. Sample heat inactivation is a key health and safety requirement, but this causes DNA to degrade (14) and template of sufficient quantity and quality for nanopore sequencing is rarely recovered. Furthermore, the low proportion of Mycobacteria DNA in sputum, eg 0.01% (15) leads to poor genome coverage eg 0.002 – 0.7X by Illumina (16).

Methods published to date for direct from sputum sequencing of MTB are relatively complex and have not been widely adopted. MTB enrichment using SureSelect hybridisation and amplification (Agilent, USA) yielded 90% to complete genome assemblies, allowing antimicrobial susceptibility prediction (17, 18) and informing treatment for one patient in real-time (19). An alternative approach using kit-based depletion of non-target DNAs (16) obtained wide variation in genome coverage (<12% to >90%). Both approaches included heat inactivation, the former at 80°C for 50 min and the latter at 95°C for 30 min (15, 17). These methods depend on commercial kits which inflate cost, shipping/storage requirements, and protocol complexity.

Our aim was to develop a simple, robust, low-cost method of preparing DNA of sufficient quality for nanopore sequencing, directly from positive sputum samples. Heat inactivation was essential, but culture and DNA amplification were excluded. The method was to be immediately transferable to two high burden settings for field testing – India and Madagascar.

## METHODS

### Research Ethics Statement

The protocol for this study was approved by London – Queen Square Research Ethics Committee (17/LO/1420). Human samples were collected under approval of East Midlands Research Ethics Committee (08/H0406/189) and all subjects gave written informed consent in accordance with the Declaration of Helsinki.

### Mock Clinical Samples for Method Development

A model system comprising standardised mock clinical samples was established by pooling infection negative human sputum samples and spiking with enumerated *Mycobacterium bovis* (BCG) Pasteur strain at known concentrations.

#### (i) Culture and Enumeration of BCG Cells

Culture conditions for BCG cells were optimised to yield mostly single cells which could be stained and counted, rather than rafts comprising large numbers of cells. Freshly prepared Bactec Mycobacteria growth indicator tube (MGIT) (Becton Dickinson, Wokingham, United Kingdom), UK) were inoculated very sparsely with 10 μl BCG Pasteur frozen stock. After 30 days incubation at 37°C, the culture was vortexed vigorously. Larger clusters of BCG cells were allowed to settle for 10 min. Fresh MGIT tubes containing 0.5% Tween 80 (Acros Organics, Geel, Belgium), to encourage non-clustered cell growth (20), were inoculated using 200 μl of the ‘settled’ BCG culture. The tubes were incubated for 18 days incubation at 37°C, then BCG cells were harvested and counted. After vigorous vortexing, 1 ml fluid was removed and BCG cells were pelleted by centrifugation for 10 min (13,000 rpm). The pellet was resuspended in 100 μl crystal violet stain (Pro Lab Diagnostics, Birkenhead, UK). Cells were counted using a Petroff Hausser counting chamber (Hausser Scientific, Horsham, PA, USA) for bacteria enumeration. The enumerated BCG stock was stored at −20°C in 1 ml aliquots until required.

#### (ii) Combining Enumerated BCG Cells and Infection Negative Sputum

Negative human sputum samples were obtained anonymously from asthmatic patients (see research ethics statement). Up to ten samples were pooled, then liquefied by treatment with an equal volume of 2x strength thermo-protection buffer (4 M KCl, 0.05 M HEPES buffer pH 7.5 (Sigma Aldrich, MO, USA), 0.1% DTT (Roche, UK), nuclease free molecular biology grade water) to ensure a final concentration of 2 M KCl. Fresh buffer was made weekly and stored in the dark at 4°C. Sputum was incubated at 37°C with occasional vortexing, until liquefaction was complete. The enumerated BCG cell stock was thawed, and a 10-fold dilution series was made in PBS, from 10^5^ to 10^1^ cells per 200 μl. The dilution series was spiked into 800 μl aliquots of the liquefied sputum in 2 ml screw cap tubes to make 1ml mock clinical samples. Microscopy was performed on these mock samples using ZN staining, and GeneXpert semi quantitative, cartridge based PCR (Cepheid, Solna, Sweden) for MTB/RIF Ultra according to the manufacturer’s instructions.

### Validation of Mycobacteria Heat Inactivation

A validation experiment was performed which confirmed that viable Mycobacteria did not survive heat inactivation at 99°C for 30 min in thermo-protection buffer (Table 1). This ‘heat-kill’ validation was performed prior to using the method on *Mycobacterium tuberculosis* positive clinical samples or MGIT cultures. Identical control samples prepared in parallel were incubated for 30 min at room temperature (Table 1). To assess Mycobacteria viability post-heating, each sample was added to a freshly prepared MGIT tube. These were checked regularly for Mycobacteria growth during incubation at 37°C for 8 weeks (or until positive). Löwenstein-Jensen slopes were also inoculated for the heat-treated samples.

**Table 1.** Heat Inactivation Validation

| Sample type | Mycobacteria cells*<sup>1</sup> | Heat treatment | Time to positive MGIT culture |
| --- | --- | --- | --- |
| (i) 1 ml pooled negative human sputum liquefied with thermo-protection buffer containing DTT. | MTB H37Rv | Room temp 30 min (control) | 10 days |
|  | MTB H37Rv | 99°C 30 min | Negative |
|  | BCG Pasteur | Room temp 30 min (control) | 18 days |
|  | BCG Pasteur | 99°C 30 min | Negative |
| (ii) 1 ml sputasol treated sputum*<sup>2</sup> to which an equal volume of thermo-protection buffer was added | MTB H37Rv | Room temp 30 min (control) | 7 days |
|  | MTB H37Rv | 99°C 30 min | Negative |
|  | BCG Pasteur | Room temp 30 min (control) | 8 days |
|  | BCG Pasteur | 99°C 30 min | Negative |
| (iii) 0.5 ml positive MGIT culture plus equal volume thermo-protection buffer | MTB H37Rv | Room temp 30 min (control) | 2 days |
|  | MTB H37Rv | 99°C 30 min*<sup>3</sup> | Negative |
|  | BCG Pasteur | Room temp 30 min (control) | 3 days |
|  | BCG Pasteur | 99°C 30 min*<sup>3</sup> | Negative |
| (iv) 1 ml positive MGIT culture spun down. Pellet resuspended in 1 ml thermo-protection buffer | MTB H37Rv | Room temp 30 min (control) | 4 days |
|  | MTB H37Rv | 99°C 30 min | Negative |
|  | BCG Pasteur | Room temp 30 min (control) | 3 days |
|  | BCG Pasteur | 99°C 30 min | Negative |

The final concentration of KCl used in each heat inactivation experiment (i) to (iv) was 2M.
\*<sup>1</sup>Spiking inoculum for sputum comprised live cultured BCG or *M. tuberculosis* H37Rv cells prepared by pelleting cells from 1 ml MGIT culture by centrifugation at 13,000 rpm for 10 minutes, then resuspending in PBS (1 ml). One drop was used as the inoculum.
\*<sup>2</sup>Sputum samples received by the Clinical Microbiology Laboratory, John Radcliffe Hospital, Oxford, without a request for TB testing were decontaminated by treatment with 4% NaOH (E and O Laboratories Ltd, Bonnybridge, Scotland), neutralised, spun down, and resuspended in 1ml sputasol. They were then spiked with 1 drop inoculum\*<sup>1</sup>.
\*<sup>3</sup>A precipitate formed on heating with thermo-protection buffer, possibly comprising salt/antibiotics/media components.

### Clinical Samples

Mycobacteria-positive clinical respiratory samples comprised sputum (n=16), bronchoalveolar lavage (BAL) (n=1) and lymph node biopsies (n=3) (the latter underwent ‘beating’ with large glass beads in saline solution for routine diagnostic testing prior to receipt). Samples were submitted for routine testing at Birmingham Heartlands Hospital NHS Foundation Trust, Birmingham, United Kingdom (n=6), or the Clinical Microbiology Laboratory, John Radcliffe Hospital, Oxford University NHS Foundation Trust, Oxford (n=14). Oxford samples had treatment with an equal volume of Sputasol (Oxoid Limited, Basingstoke, UK) prior to receipt, and were stored at 4°C. Samples from Birmingham comprised untreated sputum shipped overnight on ice to Oxford, after which they were stored at 4°C. Prior to receipt, Microscopy (with auramine staining), had yielded acid fast bacilli scores of +1 to +3 and, or a positive MTB/RIF Ultra GeneXpert result (Cepheid, Solna, Sweden). Samples were used only after routine diagnostic tests had been completed, therefore sample quality (available volume, storage time etc) varied. Samples were processed as soon as possible after receipt.

### Clinical Sample Liquefaction and Heat Inactivation

All available clinical sample volume was used. Samples were liquefied using an equal volume of thermo-protection buffer containing DTT. 1 ml aliquots were heat-inactivated in 2 ml screw cap tubes at 99°C for 30 min. Cells were collected by centrifugation (6,000 x g, 3 min) and the supernatant discarded. Then cell pellets were combined (if >1 available per sample) in a total volume of 1 ml PBS. Cells were again collected by centrifugation (6,000 x g, 3 min) and resuspended in PBS followed by another centrifugation step. The two wash steps aimed to reduce contamination with non-target DNA. The final cell pellet was resuspended in 100 μl PBS, then total DNA was extracted.

### Total DNA Extraction

0.08-0.1 g silica beads (Lysing Matrix B, MP Biomedicals, CA, USA) were added to the heat inactivated cell suspension, which underwent two rounds of mechanical disruption using an MPBio Fast Prep-24 machine (MP Biomedicals, CA, USA) at 6.0 m/s for 40 s (5 min interval). After centrifugation at 16,000 x g for 10 min at room temperature, up to 100 μl of supernatant was transferred to a fresh tube (1.5 ml DNA LoBind, Eppendorf, Hamburg, Germany). DNA in the supernatant was purified using Agencourt AMPure XP beads (Beckman Coulter, CA, USA). An equal volume of beads was added to the supernatant and incubated on a hula mixer at room temperature for 10 min. Beads with DNA bound were magnetically separated and the supernatant was removed when clear. The beads were washed using 200 μl freshly prepared 70% ethanol which was removed after a 20 s incubation. This step was repeated once more, removing as much supernatant as possible at the end of the incubation and air drying for <1 min. DNA was eluted from the beads in 50 μl 1x TE buffer (pH 8, Sigma Aldrich, MO, USA) at 35°C for 10 min. DNA concentration was measured by Qubit Fluorometer (Invitrogen, CA, USA) and the DNA Integrity Number (DIN) and fragment size by TapeStation (Agilent, CA, USA).

### ONT Library Preparation and Sequencing

Undigested DNA (up to 90 ng) was prepared for ONT sequencing using the Ligation Sequencing kit (SQK-LSK109). When samples were run multiplexed (more than one per flow cell), the Native Barcoding Expansion kit (EXP-NBD104) was used. The manufacturer’s protocols ‘Genomic DNA by Ligation’ and ‘Native barcoding genomic DNA’ were followed with minor amendments; 0.8 x volume of AMPure XP beads were used to purify the end-prep and barcode ligation reactions, incubation time with AMPure XP beads was doubled, and elution was performed at 35°C. Multiplexed sequencing libraries comprised 6 barcoded DNA samples and all libraries were sequenced using R9.4.1 SpotON flow cells on GridIONs with the MinKNOW and Guppy software versions current at the time of sequencing.

### Bioinformatics

Nanopore reads were basecalled using Guppy (Oxford Nanopore Technology, Oxford, UK). When one sample was sequenced per flow cell (without multiplexing), all the reads in the sequence data were analysed. For multiplexed runs with more than one sample per flow cell, we used Porechop (v0.2.2, https://github.com/rrwick/Porechop) to perform stringent barcode demultiplexing to minimize the number of misclassified reads. Porechop searches for the presence of the barcode sequence at both the start and end of each read; reads were classified only if the same barcode was found at both ends, otherwise the read was discarded. This level of stringency was achieved by setting the “require_two_barcodes” option in Porechop and setting the threshold for the barcode score at 60. (Porechop was used because much of the sequencing was performed prior to the availability of deepbinner or guppy_barcoder).

To allow the correct identification of *Mycobacterium tuberculosis* complex (MTB complex) reads from the sequencing data, we used both taxonomical classification and mapping approaches. Firstly, reads from each sample were taxonomically classified against the refSeq database using Centrifuge v1.0.3 (21). A read was considered as candidate for MTB complex if it was uniquely assigned to a species within MTB complex or equally assigned to more than one species within MTB complex. Human reads were discarded and not retained as part of our in house CRuMPIT workflow (22). Then, reads were mapped to either BCG (GenBank AM408590.1; the 16S rRNA region {1498360, 1499896} was masked) or TB (NC_000962.2; the 16S rRNA region {1471846, 1473382} was masked) reference sequences using Minimap2, (23) depending on the type of the sample. Reads were retained if more than 85% of the bases were mapped (ie if the length of a read is 1,000 bp, >850 bp were required to be mapped to the reference sequence). Finally, MTB complex reads were identified as those agreed by Centrifuge and mapping. Integrative Genomics Viewer was used to view the mapping profiles (24). The mapping coverage and depth across the whole genome and 22 genes associated with susceptibility/resistance to clinically important antimicrobials (25) were analysed using Samtools and Pysam (https://github.com/pysam-developers/pysam).

## RESULTS

Our initial experiments focused on identifying a buffer in which Mycobacteria DNA was protected from degradation during heat-inactivation. Living organisms can survive at temperatures around the boiling point of water (26), indicating that DNA can exist intact at high temperatures. The high concentrations of KCl and MgCl_2_ found in some hyperthermophiles are thought to help protect their DNA against thermo-degradation. This has been reproduced *in vitro* using plasmid DNA (26, 27) and formed the basis of buffer optimisation experiments.

### Optimisation of DNA thermo-protection buffer composition and heating duration

Three different 118 ng DNA extracts were made: (i) BCG DNA, (ii) BCG and sputum DNA, (iii) sputum DNA. These were heated at 99°C for 30 min (Oxford Clinical Microbiology Laboratory health and safety requirement) in four different buffers; 25 mM HEPES pH 7.5 plus 0, 0.5, 1 or 2 M KCl, then DNA (ng) remaining post-heating was recorded (Fig. 1A). The mass of DNA post-heating increased with increasing KCl (Fig. 1A). Furthermore, BCG DNA was better protected than sputum DNA; at 2 M KCl minimal BCG DNA degradation occurred, while >50% of sputum DNA degraded (Fig. 1A), indicating potential for BCG enrichment.

**Fig. 1.**
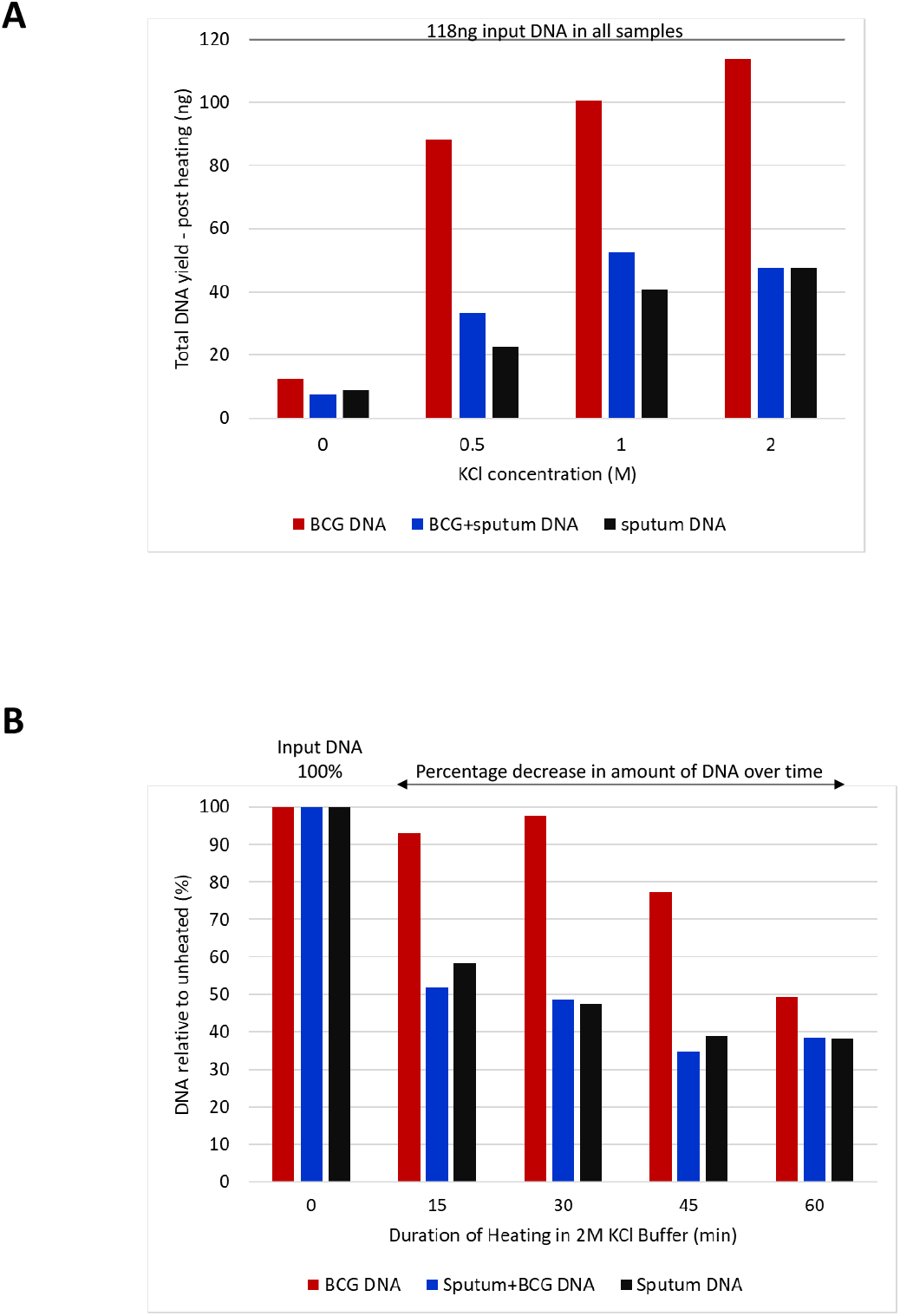
Optimisation of DNA thermo-protection buffer composition and duration of heat inactivation at 99°C. (A) Extracted DNA was heated in 25 mM HEPES buffer pH 7.5 containing 0, 0.5, 1 or 2 M KCl. Input DNA comprised 118 ng of (i) BCG DNA, (ii) BCG and sputum DNA, (iii) sputum DNA. Each DNA type was heated at 99°C, for 30 min. (B) Impact of heating duration on DNA yield. DNA remaining post heating is expressed as a percentage of the input DNA for (i) 10^5^ BCG cells, (ii) 1ml sputum spiked with 10^5^ BCG cells, or (iii) 1ml sputum. BCG DNA degraded more slowly than sputum DNA, indicating the potential for enrichment relative to human DNA at earlier time points.

Next, we determined the impact of heating duration (0, 15, 30, 45, 60 min at 99°C) on the same three extracted DNAs, in 2M KCl thermo-protection buffer. The percentage decrease in amount of DNA remaining after heating was plotted relative to input DNA (Fig. 1B). DNA yield declined over time, but BCG DNA was again more heat stable than sputum DNA. The 30 min time point was identified as ideal for both BCG enrichment and met health and safety requirements.

### Thermo-protection of DNA in intact BCG cells

Next, we investigated whether DNA within intact Mycobacteria cells could be protected by thermo-protection buffer. BCG cells (10^5^ in total) were suspended in 1 ml thermo-protection buffer and incubated at 99°C for 0, 15, 30, 45 or 60 min. Control cells were heated for the same times in phosphate buffered saline (PBS). The experiment was performed in triplicate, then the DNA was extracted. The DNA yield from BCG cells heated in thermo-protection buffer was markedly higher than those heated in PBS (Fig. 2A) except at time 0 without heating. Here, the yield of DNA was lower than expected because the cell pellet was more diffuse when it had not been heated, and cells were more easily lost than in PBS.

**Fig. 2.**
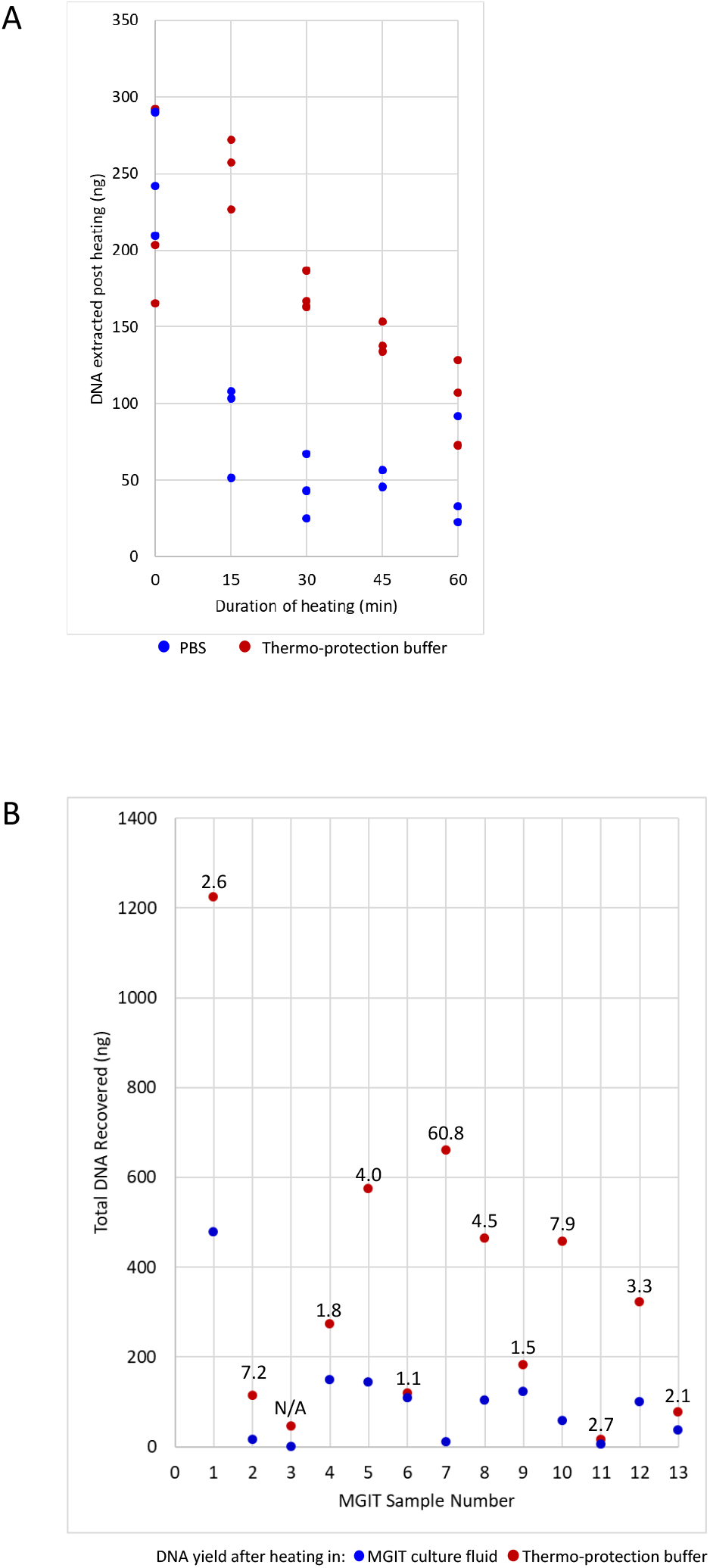
Thermo-protection of DNA in intact BCG cells. (A) Effect on DNA yield of heating intact enumerated ‘de-clumped’ BCG cells in thermoprotection buffer for the times shown. 10^5^ BCG cells were heated at 99°C for 0, 15, 30, 45 or 60 min in 2 M KCl and 25 mM HEPES pH 7.5 (thermo-protection buffer) or PBS (control). The experiment was performed in triplicate and DNA was extracted post heating. (B) DNA yield obtained when heating intact Mycobacteria cells from positive MGIT culture in thermo-protection buffer versus heating in MGIT culture fluid. Data are shown for 13 positive MGIT cultures. The DNA yield obtained after heating for 30 min at 99°C in thermo-protection buffer, compared to heating in MGIT fluid is plotted. Each dot indicates the total DNA recovered (ng) from 1 ml initial MGIT culture. Numbers above the dots indicate the fold improvement in DNA yield when thermo-protection buffer was used rather than MGIT culture fluid. N/A indicates a sample where no DNA was recovered after heating in MGIT culture fluid, so no ‘fold improvement’ could be calculated.

In a separate experiment using intact Mycobacteria cells, 13 positive MGIT cultures (anonymised discards obtained from Oxford Clinical Microbiology Laboratory) were heated in 1ml thermo-protection buffer or in culture fluid. DNA yield was improved for the cells heated in thermo-protection buffer (Fig. 2B).

### Confirmation of Mycobacteria Heat Inactivation

A validation experiment was performed to confirm that viable Mycobacteria (MTB H37Rv or BCG Pasteur) did not survive 30 min heating at 99°C in thermo-protection buffer. After 8 weeks incubation at 37°C no growth occurred in the heated samples. In contrast, the room temperature controls remained viable (Table 1).

### Direct-from-Sample Sequencing of BCG-spiked Mock Clinical Samples

Four sets of mock clinical samples were made, each set containing a ten-fold dilution series of enumerated BCG cells (10^5^ – 10^1^ and zero cells) in 1 ml infection negative human sputum (liquefied in thermo-protection buffer). Four different batches of pooled sputum were used, but the BCG cells were from the same enumerated batch. All four replicates (experiments A-D) underwent heat inactivation (99°C for 30 min), DNA extraction and ONT sequencing using a single flow cell per sample. Replicates in experiments B, C and D underwent additional microscopy (ZN staining) and GeneXpert PCR.

Sequencing, Microscopy and GeneXpert PCR yielded reproducible data across the replicate experiments (Fig. 3, Table 2). The number of MTB complex sequencing reads generated per sample was linear and indicated detection down to 10^1^ input BCG cells. At this concentration, 31, 49, 51, and 59 MTB complex reads were detected (Fig. 3A). At 10^3^ BCG cells input, genome coverage (1x) was >90% (Table 2). The ratio of human reads to MTB complex reads was also linear and reproducible (Fig. 3B).

**Fig. 3.**
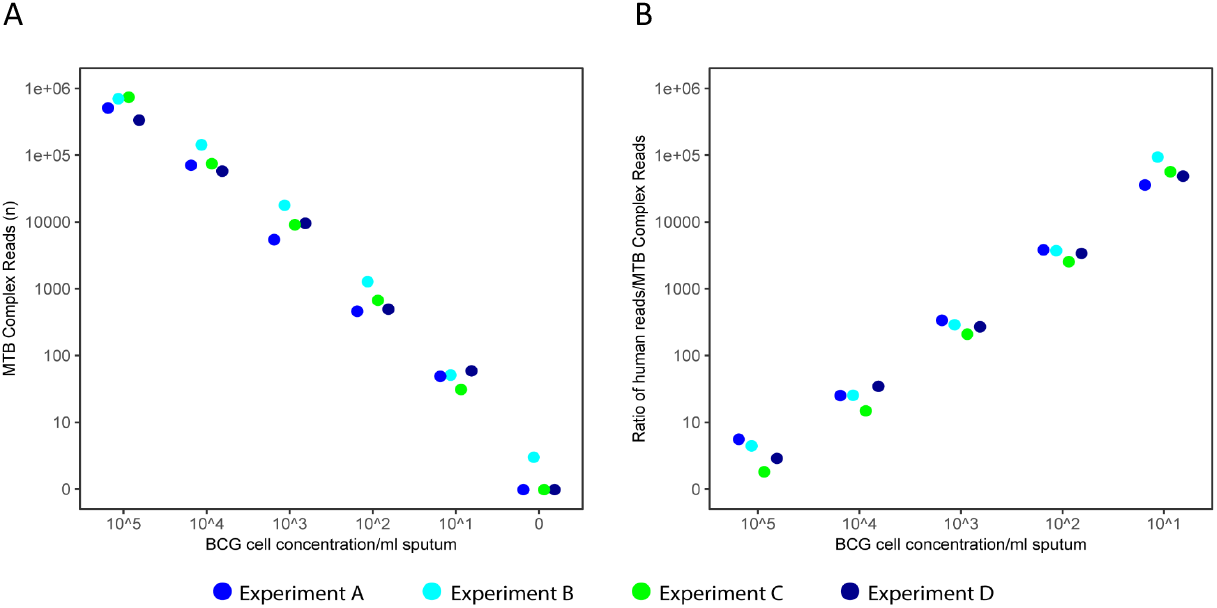
Validation of DNA thermo-protection method using Mock Clinical Samples. Mock clinical samples containing enumerated BCG cells (0 – 10^5^) in 1ml infection negative human sputum liquefied in thermo-protection buffer underwent heat-inactivation at 99°C for 30 min. DNA was extracted and sequenced on a ONT MinION (1 R9.4.1 flow cell per sample). Reproducibility was assessed using four replicate experiments (A-D). (A) Number of MTB complex reads generated per sample was linear and indicated a detection limit of 10^1^ BCG cells. (B) Ratio of human reads to MTB complex reads.

**Table 2.** Reproducibility and detection limits of Microscopy, GeneXpert, and direct from sample ONT Sequencing.

| BCG cells | Microscopy | GeneXpert |  | Nanopore |  |  |  |
| --- | --- | --- | --- | --- | --- | --- | --- |
| Experiment B | ZN Stain | Ct value | Detection | MTB complex Reads (n) | Mean Depth | Genome covered 1x (%) | Genome covered 5x (%) |
| 10<sup>5</sup> | +++ | 16.4 | High | 701,436 | 261.99 | 98.73 | 98.12 |
| 10<sup>4</sup> | +++ | 16.5 | High | 142,918 | 55.0 | 98.30 | 97.75 |
| 10<sup>3</sup> | + | 17.1 | Medium | 17,798 | 7.17 | 97.86 | 83.56 |
| 10<sup>2</sup> | (+) | 19.3 | Low | 1,280 | 0.56 | 44.26 | 0.04 |
| 10<sup>1</sup> | - | 23.0 | Very Low | 51 | 0.03 | 2.92 | 0 |
| 0 | - | - | Negative | 3 | 0 | 0.06 | 0 |
| **Experiment C** |  |  |  |  |  |  |  |
| 10<sup>5</sup> | +++ | 16.2 | High | 738,605 | 225.73 | 98.48 | 97.91 |
| 10<sup>4</sup> | +++ | 16.3 | High | 74,731 | 20.19 | 97.76 | 97.12 |
| 10<sup>3</sup> | + | 17.1 | Medium | 9,086 | 2.75 | 91.84 | 17.98 |
| 10<sup>2</sup> | + | 17.6 | Medium | 672 | 0.21 | 19.11 | 0 |
| 10<sup>1</sup> | - | 22.7 | Low | 31 | 0.01 | 1.38 | 0 |
| 0 | - | - | Negative | 0 | 0 | 0 | 0 |
| **Experiment D** |  |  |  |  |  |  |  |
| 10<sup>5</sup> | +++ | 16.2 | High | 333,937 | 99.92 | 98.15 | 97.62 |
| 10<sup>4</sup> | ++ | 16.4 | High | 57,824 | 16.28 | 97.66 | 97.13 |
| 10<sup>3</sup> | + | 17.0 | Medium | 9,533 | 3.37 | 93.97 | 29.65 |
| 10<sup>2</sup> | + | 18.9 | Low | 494 | 0.19 | 17.51 | 0 |
| 10<sup>1</sup> | - | 23.0 | Very Low | 59 | 0.02 | 2.29 | 0 |
| 0 | - | - | Negative | 0 | 0 | 0 | 0 |

Bioinformatics methods were optimised to ensure reads in negative controls (such as rRNA genes from non-target bacterial species (28)) were not incorrectly assigned as BCG; prior to these improvements close to 10,000 reads were incorrectly identified as MTB complex in the negative control (Fig. S1A). After the improvements, the negative controls for experiments A, C and D contained zero MTB complex reads, however three Mycobacteria reads were present in the negative control of experiment B (Fig. 3A, Figure S1A, Table 2).

GeneXpert (Cepheid) and microscopy results also followed the concentration of BCG cells spiked into each sample (Table 2). The detection limit of GeneXpert was 10^1^ BCG cells and microscopy 10^2^ where cells were described as very scanty, ie 1 or 2 per 100 fields (Table 2).

### Direct from Sample Sequencing using Multiplexing

Sequencing more than one sample per flow cell (multiplexing), offers both time and cost efficiencies. To assess its feasibility, a short DNA ‘barcode’ was ligated to each DNA sample, then the 24 DNAs from replicate experiments A to D were sequenced at six samples per flow cell – one per replicate experiment. After sequencing, the barcodes were identified bioinformatically and the data were assigned to their original sample. Unfortunately, the 10^1^ and 10^2^ BCG spiked samples contained a similar number of MTB reads to the negative control (Fig. S1B), therefore using this approach the limit of detection declined 100 fold to 10^3^ BCG cells/ml sputum. This was a result of the barcodes of the BCG positive samples being incorrectly (and unavoidably) identified as that of the negative control (Fig. S1B). The multiplexing approach was also compromised by a reduction in the total data available for analysis. Although we applied stringent barcode demultiplexing criteria, between 5.28 and 46.9% of total reads cannot be reliably assigned to an input sample.

### Thermo-protection method enriches mock clinical samples for MTB DNA

The multiplexed data for the mock clinical samples in experiments A to D were used separately, to confirm whether the heat inactivation in thermo-protection buffer, enriched the samples for BCG sequences by depleting human DNA. This confirmed that heated samples were enriched for MTB DNA/depleted for human DNA (Fig. 4) when compared to equivalent controls prepared without heat inactivation and washing steps.

**Fig. 4.**
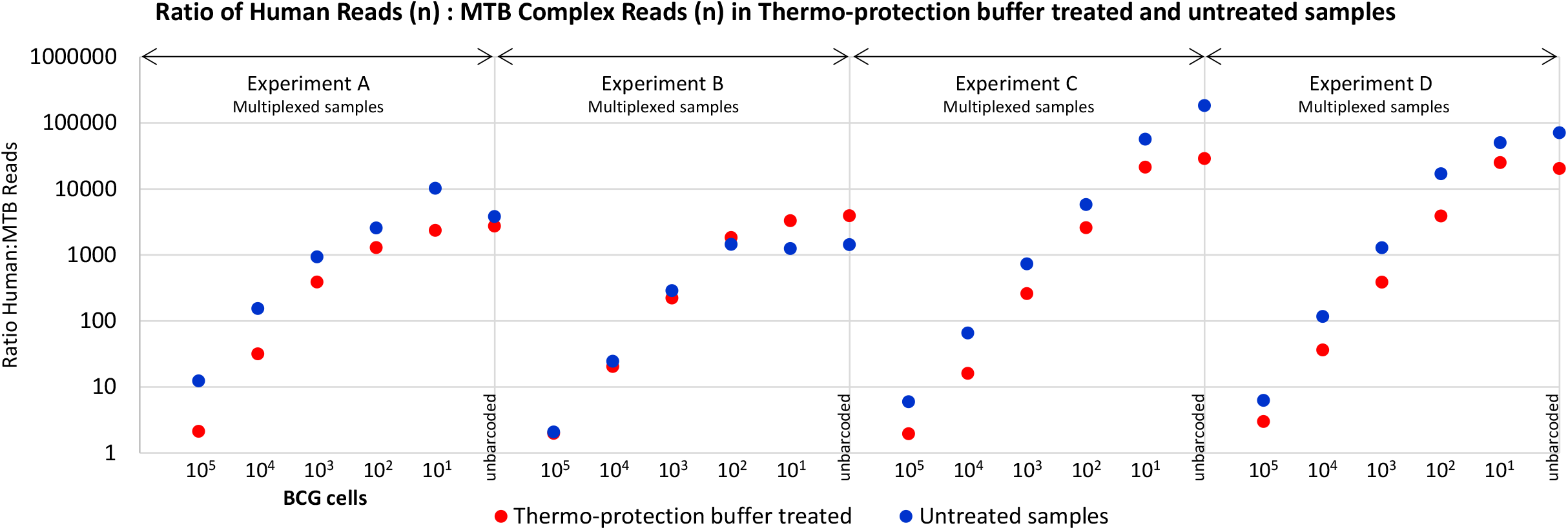
Mock clinical samples were enriched for Mycobacteria DNA after heating in thermo-protection buffer. Data are shown for four replicate experiments, A to D, in which samples were barcoded and run multiplexed six per flow cell. Each experiment comprised samples made from a batch of infection negative sputum (liquefied using thermo-protection buffer containing DTT), 1 ml aliquots of which were spiked with enumerated BCG cells at 10^5^ to 10^1^ cells, and zero BCG cells (control). Sputum batch and therefore ‘background’ DNA did not vary within replicates A to D, only between them. The full set of replicates was set up twice with heating (99°C, 30 min), and without heating. After sequencing, the numbers of BCG and human derived reads was assessed and their ratio in each sample calculated. Higher ratios of human: MTB reads were obtained for samples which were not heated in thermoprotection buffer, indicating heated samples were enriched for MTB reads relative to human reads ie human DNA was depleted. The exception to this was experiment B, which yielded anomalous results because the number of reads for the unheated sample was unusually poor.

### Direct-from-Sample Sequencing of *M. tuberculosis* Positive Clinical Samples

#### (i) DNA Preparation using thermo-protection method

A total of 20 MTB positive clinical samples comprised 16 sputa, three lymph node biopsies, and one bronchoalveolar lavage sample. Samples were 1–14 days old, and volumes ranged from 0.25 – 1.5 ml. Microscopy and GeneXpert results indicated variable MTB loads (Table 3). The total DNA extracted ranged from 105 – 3970 ng per sample, the DNA integrity number (DIN) from 1.8 – 6.3 and the peak fragment length 1,834 – 13,949 bp (Table 3). Each sample underwent direct-from-sample sequencing using a single R9.4.1 flow cell.

**Table 3.**
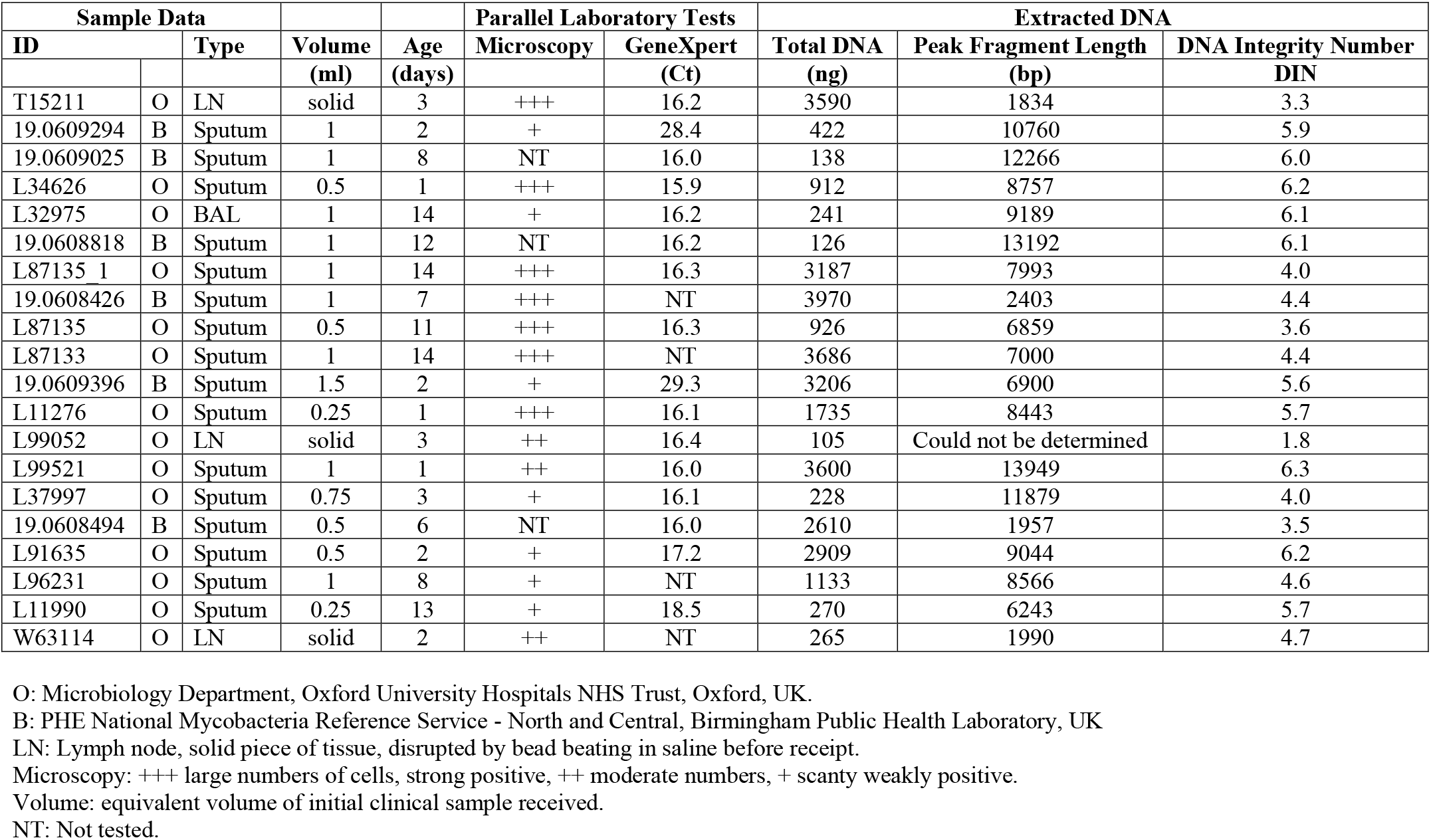
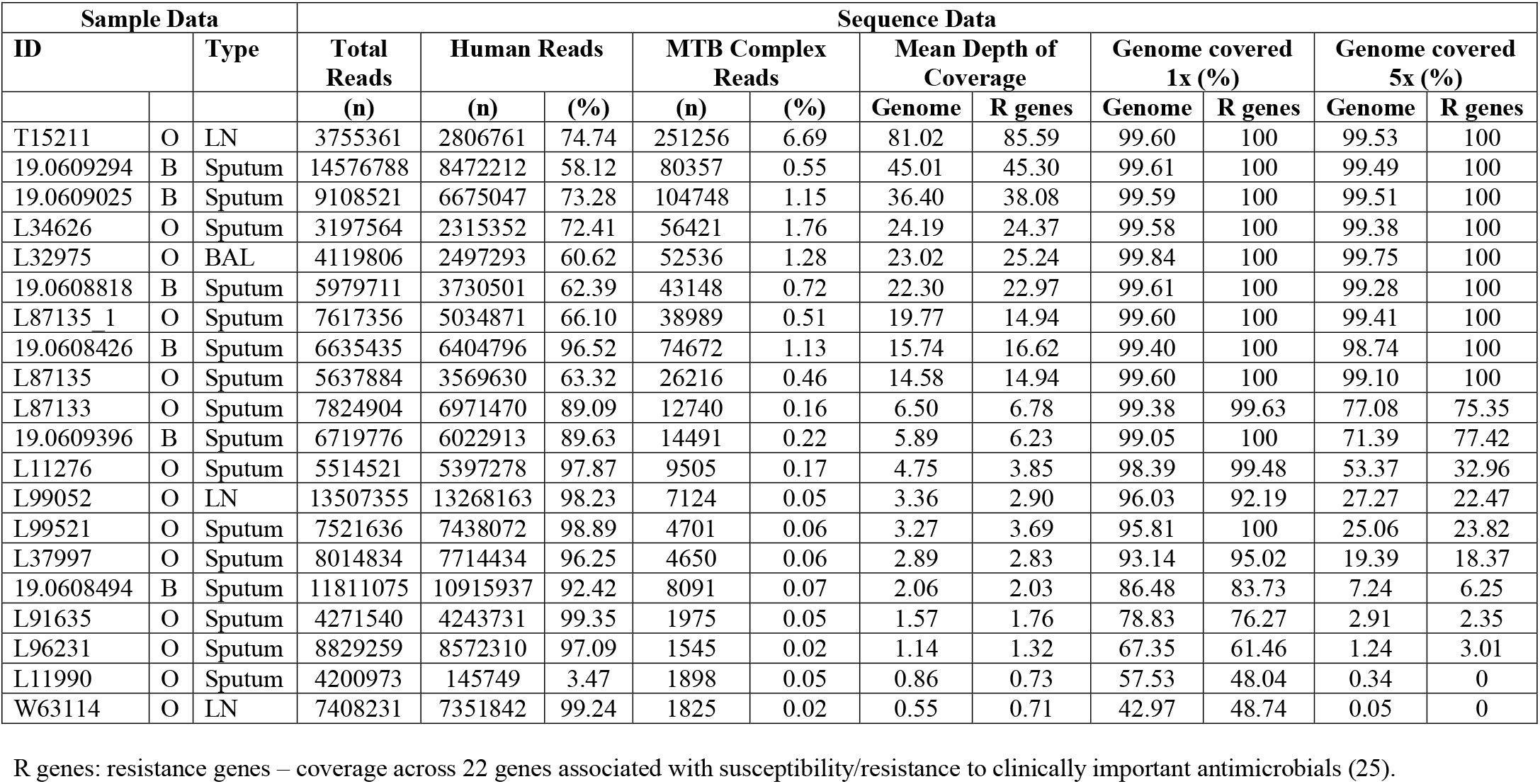
Direct ONT Sequencing of MTB Positive Clinical Samples

#### (ii) Sequence data

The total number of reads obtained per flow cell ranged from 3,197,564 to 14,576,788 (Table 3). Human reads were discarded prior to detailed analysis (ethics requirement). Among the non-human reads, mean MTB read length was up to 4.77 times longer than for non-MTB (Fig. 5A). MTB reads were detected in all twenty clinical samples (n=1,825-251,256) (Table 3), the mean depth of genome coverage ranging from 0.55 to 81.02. An initial sample volume ≥1ml, and a lower percentage of human reads was apparently associated with higher depth of coverage, although the numbers were too small for statistical analysis (Table 3).

**Fig. 5.**
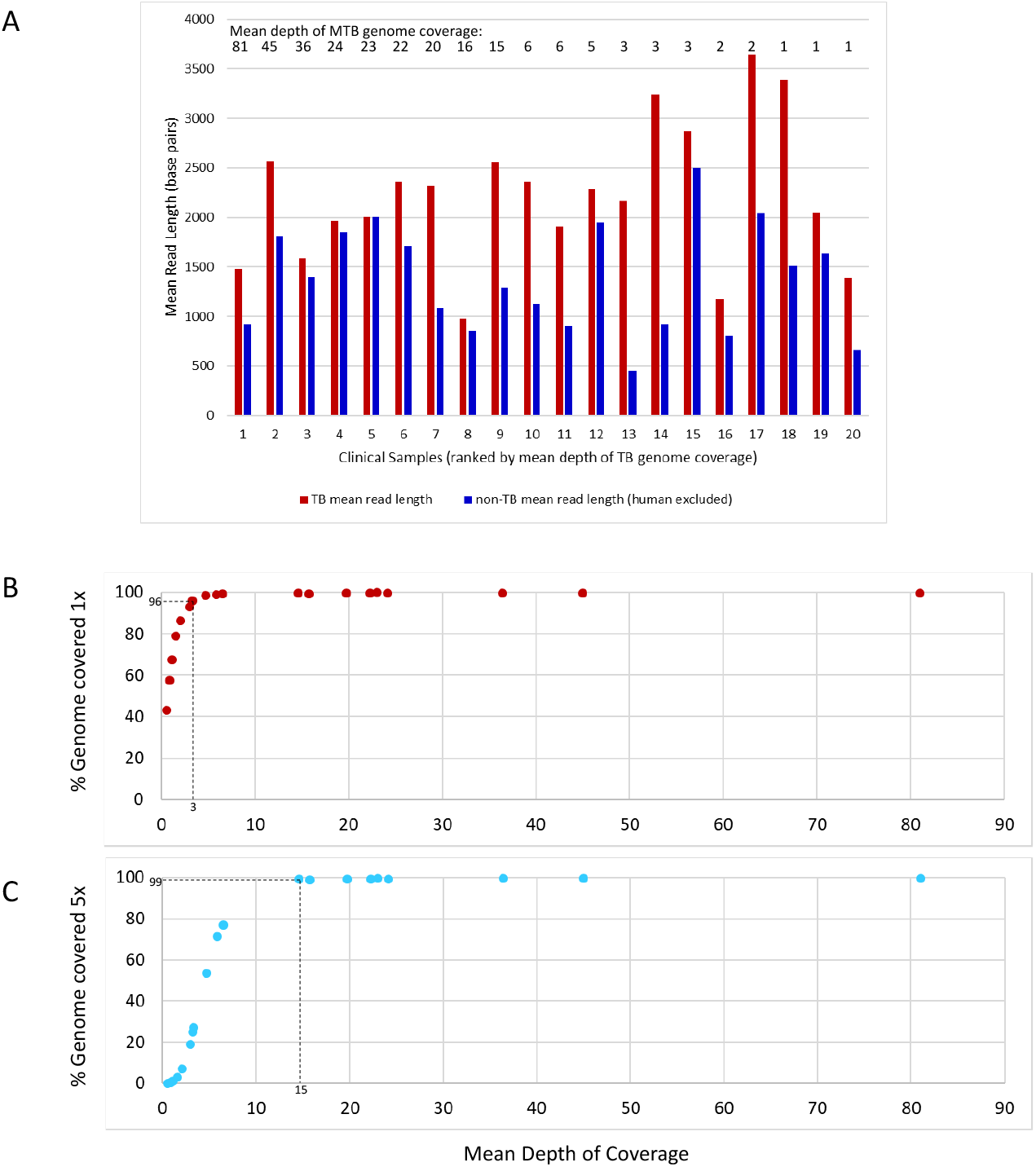
Sequence data generated direct from clinical samples – mean read lengths (MTB vs non MTB non-human sequences) and relationship between mean depth of coverage and complete genome coverage. (A) Comparison, for each of 20 clinical samples, of mean read length for MTB and non-MTB sequences (human sequences excluded prior to analysis). Clinical samples were ranked according to mean depth of coverage, indicated by numbers above the bars. (B) Relationship between mean depth of coverage and percentage of the MTB genome covered once. A mean depth of coverage of three is required to achieve >96% one-fold TB genome coverage, achieved in 15/20 clinical samples. (C) Relationship between mean depth of coverage and percentage of the MTB genome covered five times. A mean depth of coverage of 15 is required to achieve >99% five-fold genome coverage, achieved in 9/20 clinical samples.

Plotting the mean depth of coverage against the percentage of MTB genome covered once, or five times revealed that ≥96% of the genome was covered once when a mean depth of approximately three was reached, and ≥99% was covered five times after a mean depth of fifteen (Fig. 5B, C). The depth of coverage across 22 key genes used to predict susceptibility to clinically important antimicrobials (25) closely followed the mean genome coverage (Table 3) (except one rRNA gene which was bioinformatically masked), indicating the absence of bias in depth of coverage across these regions of the genome, and confirming potential for antimicrobial resistance prediction.

## DISCUSSION

The potential of the Nanopore platform for sequencing human and plant pathogens in varied and challenging locations is well established (10–13). It works on the principle of nanopore sequencing of DNA strands, with read lengths of several hundred to hundreds of thousands of bases obtained in real time (29, 30). The preparation of long, high quality input DNA is therefore essential. However, since DNA degrades rapidly at high temperatures (31) heat-inactivated MTB clinical samples typically yield poor quality material for sequencing. Here, we describe a simple, low cost method which overcomes this technical challenge. Sputum liquefaction and heat-inactivation were accomplished following addition of an equal volume of thermo-protection buffer (4M KCl, 0.05M HEPES buffer pH7.5, and 0.1% DTT) which inhibited DNA degradation during incubation at 99 °C. Samples were fortuitously enriched for Mycobacteria DNA under these conditions (Fig. 1, 2, 4). Buffer addition was the only handling step involving infectious material, minimising risk to staff in settings where containment laboratories are not available.

The composition of thermo-protection buffer was designed to emulate intracellular conditions of hyperthermophiles (26). At high temperatures it is thought that intracellular salts such as KCl and MgC1_2_ protect the DNA’s N-glycosidic bonds against depurination and cleavage by hydrolysis of the adjacent phosphodiester bond (14, 41, 42). We chose K^+^ over Mg^2+^ because high K^+^ concentrations protect against cleavage at apurinic sites, while high Mg^2+^ concentrations stimulate this (42). Furthermore, plasmid DNA appeared better protected in KCl (Fig. 1 in 42). The choice of KCl concentration (2 M) was informed by our own data (Fig. 1) and published data (26, 27). The mechanism whereby DNA in intact Mycobacteria cells was protected during heating in thermo-protection buffer (Fig. 2) is unclear, but suggests that the cell is, or becomes permeable to K^+^ during heating. Our data (Fig. 2B) indicate that thermo-protection buffer can also improve the DNA yield obtainable from positive MGIT cultures, as currently used clinically by Public Health England for routine Illumina sequencing (43, 44). This may help to reduce numbers of samples which fail due to a low DNA yield.

The Oxford clinical microbiology laboratory chooses to inactivate Mycobacteria positive samples at 99°C for 30 min, because less stringent conditions (eg 20 min at 80°C) show variable efficacy (32–38). We confirmed that Mycobacteria heated in thermo-protection buffer at 99°C for 30 min were not viable (Table 1). Direct from sample sequencing results are optimal when input DNA is enriched for sequences of interest (16–18, 39). Depletion of up to 99.99% human DNA from non-TB lower respiratory tract samples has been achieved using saponin, osmotic shock, and ‘high salt’ nuclease treatments (40). However, no heat-inactivation was performed, and a specialist nuclease (Salt Active Nuclease, ArcticZymes, Tromsø, Norway) was required. Interestingly, we observed that during heating in thermo-protection buffer for 30 min at 99°C, non-target DNA degraded more rapidly than Mycobacteria DNA, providing fortuitous enrichment (Fig. 1, 2, 4). Consistent with this, mean read length obtained for MTB was longer than other non-human reads (Fig. 5A). We found that sufficient non-target DNA remained in our samples to provide a useful ‘carrier’. This was particularly important in low titre MTB samples; our 10^1^ BCG limit of detection in mock clinical samples (Fig. 3A) would not have been achieved if the majority of human ‘carrier’ DNA had been removed.

Three features of Mycobacteria DNA may have contributed to its enrichment relative to human DNA (Fig. 1, 4). Firstly, Mycobacteria DNA has a higher GC content (*M. tuberculosis* 65.6% GC, versus <50% GC for ~92% of human DNA (45, 46). Secondly, intact MTB chromosomes are covalently closed circles – resistant to thermo-denaturation because the two single strands remain intertwined during heating (47). Finally, the MTB chromosome is negatively supercoiled (underwound – a feature potentially connected to its slow growth rate (48)), but less so than some bacterial species including *E.coli* (49). Lesser negative supercoiling reduces base exposure (50), which may reduce susceptibility to thermo-degradation, relative to human DNA. We did not assess the impact of heating on DNA from non-target microbes because the range and number of cells present was unpredictable and bioinformatics analysis cannot give a reliable taxonomic classification for approximately 20% of the total reads in each sample.

We obtained optimal results when using a single R9.4.1 flow cell per sample (Fig. 3, Table 3). This approach would be prohibitively expensive if used routinely, since a flow cell costs £380 – £720 depending on order size (1 to 300). Unfortunately, multiplexing six samples per flow cell did not provide a solution, since the inefficiency of barcode ligation and incorrect bar code identification post-sequencing reduced the limit of detection 100 fold (Fig. S1). Run-time flexibility is also incompatible with multiplexing, due to variations in sample MTB titres. A solution may be to wash, re-generate and re-use flow cells after each use (https://store.nanoporetech.com/flow-cell-wash-kit-r9.html), or adopt single-use ONT Flongles at a cost of £72.50 each (at 30/01/2020). Unfortunately, the latter currently offer only 60-70 active sequencing pores in our hands, compared to 1200 – 1500 pores per R9.4.1 flow cell.

The thermo-protection method was applied successfully to clinical samples (n=20). Higher mean depth of genome coverage appeared to reflect initial sample volume (≥1 ml being ideal), and a lower human DNA content, but did not necessarily correlate with microscopy or GeneXpert Ct values (Table 3). This may reflect the known variation in copy numbers of GeneXpert targeted insertion sequences (IS6110 and IS1081) between BCG (used in mock samples, where limit of detection, microscopy and GeneXpert data all correlated, Table 2) and MTB (51, 52). Also, microscopy was performed using auramine staining for clinical samples, ZN staining for mock clinical samples, and the former was performed by multiple different staff members.

The accuracy of DNA consensus sequences obtained using the Nanopore platform is 99.9% when Nanopolish is used (R.R. Wick, L.M. Judd, K.E. Holt. 2018. Comparison of Oxford Nanopore basecalling tools; https://github.com/rrwick/Basecalling-comparison#references), indicating potential for antimicrobial susceptibility prediction. Further work is required to examine this aspect in detail, particularly for ribosomal RNA genes which were masked to improve the accuracy of MTB detection, but are implicated in resistance (such as the 16s rRNS rrs gene; aminoglycoside resistance). A further potential advantage of direct from sample sequencing (using SureSelect and Illumina) is the detection of more genetic diversity than sequencing from culture (53). Increased numbers of target reads from low titre samples will require innovations such as Mycobacteria cell fractionation or concentration, followed by DNA amplification. The method could be enhanced by adapting to a cartridge-based system, further simplifying its application in resource poor settings.

In summary, a simple, low-cost method was developed to prepare MTB DNA for Nanopore sequencing direct from clinical samples. Neither commercial kits, nor time consuming culture were required, but the key health and safety requirement, heat-inactivation, was retained and exploited to achieve target sequence enrichment. Available data suggest the method can yield complete MTB genome sequences direct from clinical samples, without amplification, achieving up to 81 fold mean depth of coverage. The protocol is currently undergoing testing by collaborators in India and Madagascar, with early data indicating reproducibility.

---

**Fig. S1.**
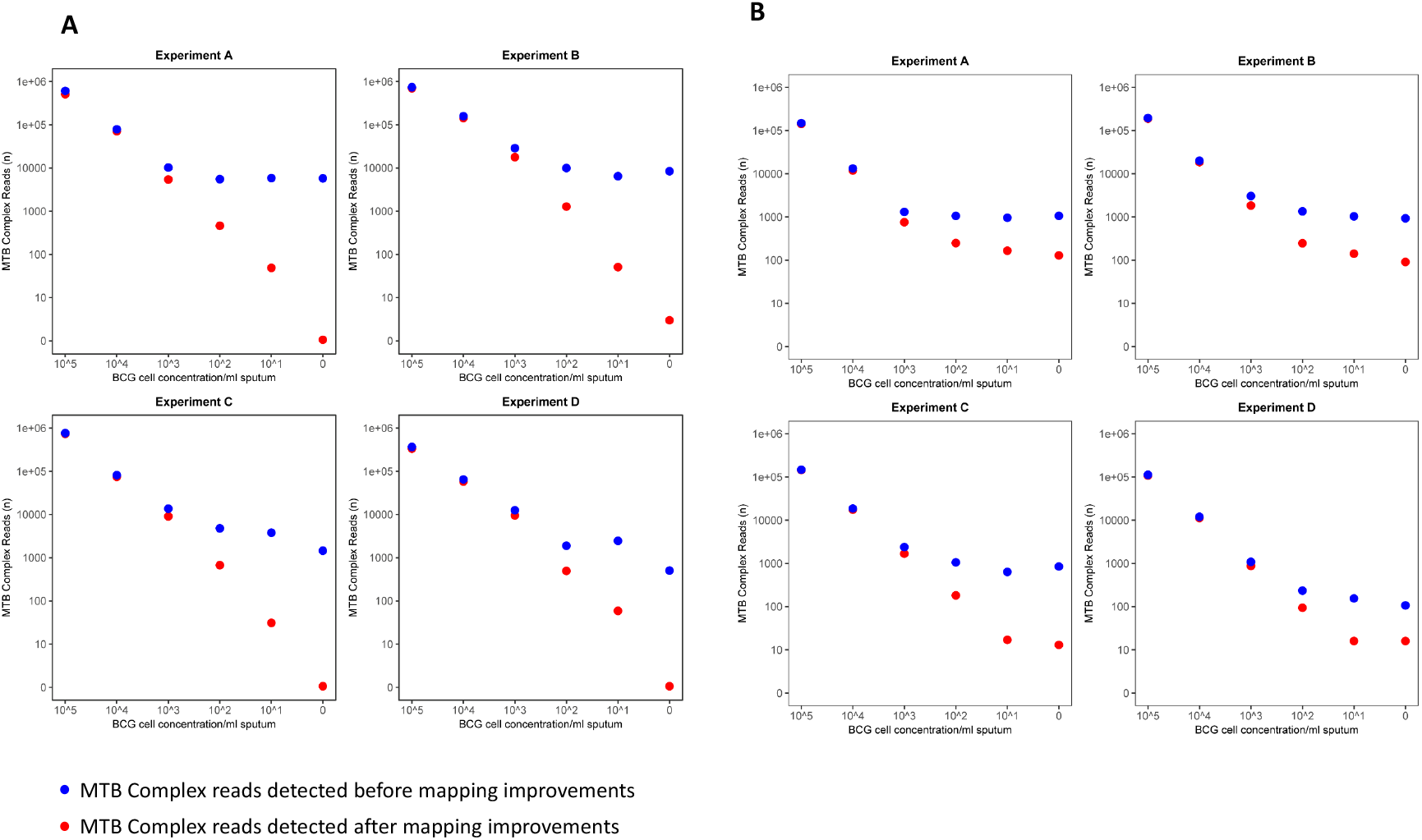
Improvements to bioinformatic analysis of Nanopore sequence data generated direct-from-sample. (A) Analysis of results generated from mock clinical samples, each sequenced on a single flow cell. Samples comprised BCG spiked infection negative sputum containing 10^5^ to 10^1^ and zero BCG cells. Data are shown before (blue dots) and after (red dots) mapping improvements. Prior to mapping improvements, close to 10,000 reads per sample were incorrectly identified as MTB complex. After the improvements, three contaminant reads were detected in the negative control (zero BGC cells) of experiment B only. These were 612, 788, and 1305 bases long and mapped independently and at high quality to the BCG reference genome, at positions 2798622-2799216, 282379-283169 and 2701067-2702371, the identity of these reads was further confirmed by BLASTn. (B) Analysis of results generated from mock clinical samples, ‘barcoded’ and sequenced multiplexed six per flow cell. The results are shown before (blue dots) and after (red dots) mapping improvements. Even after mapping improvements, many reads persisted in the low BCG titre samples and negative controls. So, when running multiplexed samples, the limit of detection was compromised at 10^3^. The reason for this was the barcodes of DNA fragments belonging to BCG positive samples were incorrectly (and unavoidably) identified as the barcode of the negative control. The multiplexing approach was also compromised by a reduction in the total data available for analysis, since a relatively high proportion of the total reads were unbarcoded.

